# Deep learning-based cell profiling based on neuronal morphology

**DOI:** 10.1101/2023.07.23.550158

**Authors:** Qiang Liu, Francesca Nicholls, Helen A. Rowland, Adrià Dangla-Valls, Shuhan Li, Yi Zhang, Piotr Kalinowski, Elena Ribe, Jamie L. Ifkovits, Sanjay Kumar, Cuong Q. Nguyen, Alejo Nevado-Holgado, Noel J. Buckley, Andrey Kormilitzin

## Abstract

Treatment of neurons with β-amyloid peptide (Aβ_1-42_) has been widely used as a model to interrogate the cellular and molecular mechanisms underlying Alzheimer’s disease, and as an assay system to identify drugs that reverse or block disease phenotype. Prior studies have largely relied on high content imaging (HCI) to extract cellular features such as neurite length or branching, but these have not offered a robust/comprehensive means of relating readout to Aβ_1-42_ concentrations. Here, we use a deep learning-based cell profiling technique to directly measure the impact of Aβ_1-42_ on primary murine cortical neurons. The deep learning model achieved approximately 80% accuracy, compared to 54% for the cell phenotypic feature-based approach. The deep learning model could distinguish subtle neuronal morphological changes induced by a range of Aβ_1-42_ concentration. When tested on a separate dataset, the accuracy remained comparable and dropped by only 2%. Our study demonstrates that deep learning-based cell profiling is superior to HCI-based feature extraction on neuronal morphology and it provides an alternative to a dose/response curve, where the modality of the response does not have to be pre-determined. Moreover, this approach could form the basis of a screening tool that can be applied to any cellular model where appropriate phenotypic markers based on genotypes and/or pathological insults are available.

## 1. Introduction

Computational cellular models, considered as a grand challenge of the 21^st^ century in systems and mathematical biology, have developed at a frenetic pace ^1 2^. Such models have successfully unveiled hidden mechanisms underlying human diseases and recapitulate characteristics of diseases at the cellular level ^3 4^, which can be deployed to accurately and rapidly identify disease phenotypes ^5 6^. Furthermore, they enable researchers to gain insights into the development of diseases and further explore potential interventions at the preclinical stage ^7^.

Alzheimer’s disease (AD) is a progressive neurodegenerative disease with nearly 7 million new cases emerging worldwide each year ^8 9^. Effective interventions are desperately desired but can only come with a fundamental understanding of the phenotypic responses to pathophysiologically relevant insults ^10^. Recent research has claimed that the production and deposition of β-amyloid peptide (Aβ_1-42_) in the human brain are correlated with AD and Mild Cognitive Impairment (MCI) ^11 12^, and can further predict early AD status ^13 14^. Ex vivo primary neurons present a powerful research model to understand neurological disease. Compared to neural cell lines, primary neurons bear molecular and cellular phenotypic features that correspond closely to their in vivo counterparts ^15^. Primary murine cortical neurons have been extensively utilized as cellular models to interrogate AD-related mechanisms. They are one of the most frequently used experimental paradigms to analyse phenotypic response to application of Aβ_1-42_, which may facilitate our understanding of the mechanism of AD and further discover new therapeutics.

Traditional image-based cell phenotyping adopts an indirect two-step approach, where a set of pre-determined subcellular compartment morphological features are first extracted and summarised from raw images ^16^. Regression or classification models are then built on these aggregated features for downstream analysis ^17^. This inevitably forces subjective human decision-making factors into the data analysis such that a significant amount of raw data that may contain subtle signals is abandoned and remain unused. Namely, the researcher has to decide in advance which subcellular features are likely to be affected by the assay. In addition, the assay may affect characteristics of the cell that cannot be compartmentalised into traditional cellular features. Machine learning (ML) has been used to classify phenotypes and has been shown to be superior to eye-based classification in several studies including classification of multiple cancer cell lines ^18^ and reversing phenotypes subsequent to genetic perturbations ^19^. More recent advances in the computer science domain, particularly the deep learning technique, deploy an end-to-end modelling approach where a model takes raw images as input and directly yields expected outcomes. In such a way, complex non-linear relationships and latent patterns uninterpretable by the naked eye can be effectively explored, and the most appropriate features relevant to the task are learned automatically, which enables them to achieve state-of-the-art performance ^20–22^. Furthermore, a model developed for one task can be easily deployed by a new one after slight parameter fine-tuning because the generic low-level features learnt by the model are transferable ^23^.

In this study, we demonstrate the development and utilization of a deep learning model to classify neuronal cell phenotype based on morphological changes in response to disease-relevant stimulus exemplified here using Aβ_1-42_ as an effective generalisable tool. Recent progress in deep learning-based methods has already demonstrated promising results in cellular image analysis across various domains ^23^. To our knowledge, this study is the first to classify cellular neurodegenerative phenotype and the results presented herein highlight the superior power of an AIML-enabled model compared to traditional two-step HCI-based cell profiling methods and human experts in terms of accuracy. This work establishes a foundation that can be further utilized in downstream tasks such as identification of small molecules or genetic interventions that block or reverse neurodegenerative phenotypes.

Additionally, the integration of deep learning methods with other advanced analytical techniques, such as single-cell sequencing or proteomics, may offer a more comprehensive view of the complex interplay between cellular components and their contribution to disease processes. By providing novel data-driven insights and facilitating the deconvolution of complex biological processes, deep learning techniques have the potential to transform our understanding of cellular behaviour and function, ultimately shaping the future of drug discovery and therapeutic development.

## 2. Results

### 2.1 Model performance

The cellular model system employed here included the primary mouse neuronal cultures that were treated with Aβ_1-42_ in 6 doses (0, 0.1, 0.3, 1, 3, 30 µM) and then immunostained with Hoechst 33342 and antibodies that recognised microtubule-associated protein 2 (MAP2), synaptophysin and postsynaptic density-95 (PSD-95). This allowed visualisation of the nucleus, neuronal cytoskeleton and localisation of pre- and post-synaptic markers. We report the accuracy of our models in Table 1. In terms of in-distribution (ID) validation of 6 doses classification, the deep neural network (DNN) model using 5-channel (MAP2, synaptophysin and PSD-95 immunostaining, Hoechst stain and bright-field) image stacks as input achieved the best performance at 80.63% accuracy. According to the confusion matrix in Figure 2, the model was capable of distinguishing neurons treated with vehicle, 0.3µM, 1µM, 3µM, and 30µM Aβ_1-42_. We anticipated that misclassifications would predominantly occur between adjacent dosages likely due to insufficient distinguishing features, and this indeed proved to be the case. Most misclassifications were made between vehicle and the lowest concentration of 0.1 µM Aβ_1-42_, indicating that the model could not accurately distinguish between these two rather close conditions. A small number of misclassifications between doses 1 and 3 µM was also observed, indicating that changes in cell morphology induced by either of these concentrations of Aβ_1-42_ were, as expected, more similar than between more distant concentrations of Aβ_1-42_. Cell morphologies of other doses were all significantly different from each other.

**Figure 1.**
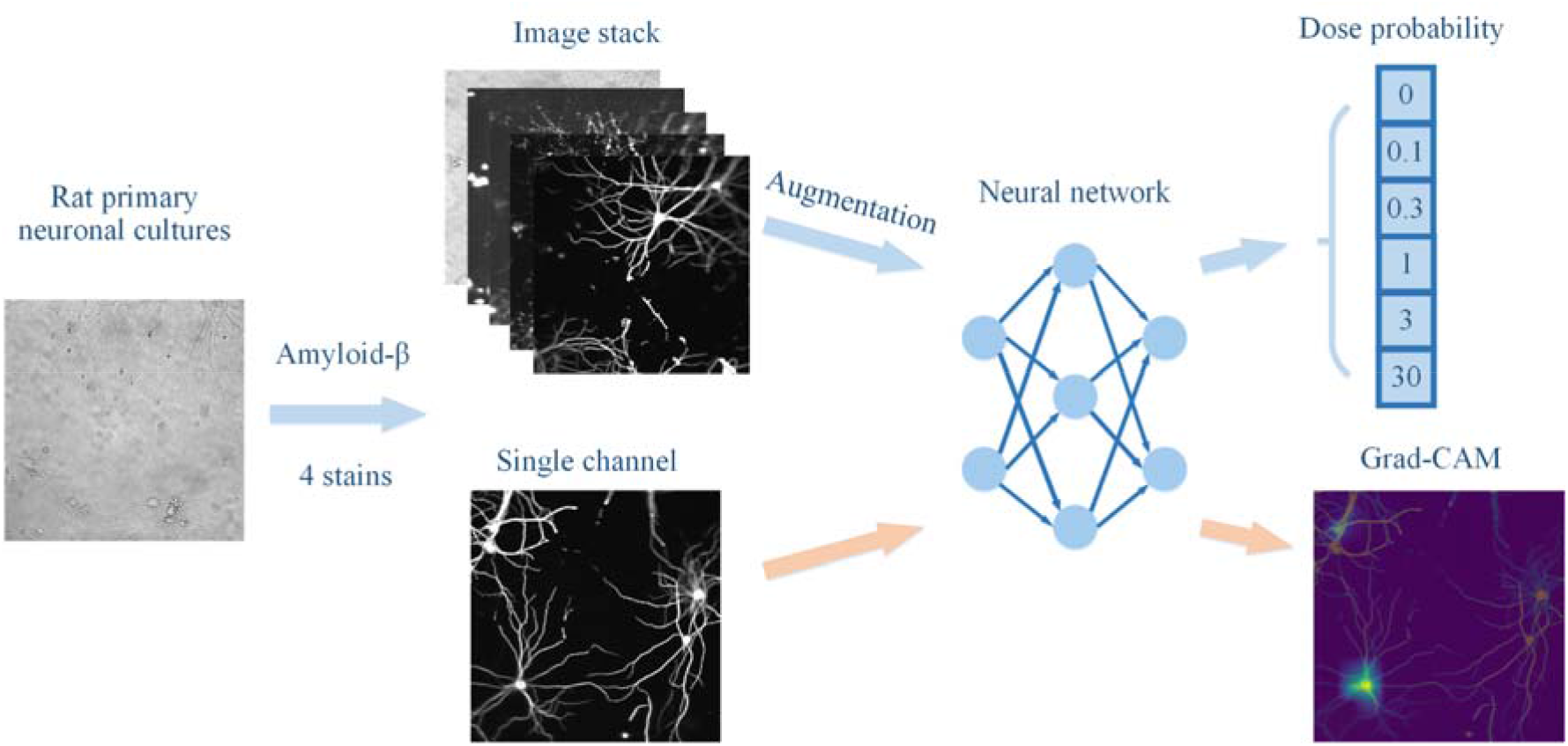
Workflow of Aβ_1-42_ treatment identification. Blue arrows: model training and classification pipelines. Orange arrows: model interpretation pipeline. Aβ_1-42_: β-amyloid peptide (1-42). Grad-CAM: Gradient-weighted Class Activation Mapping.

**Figure 2.**
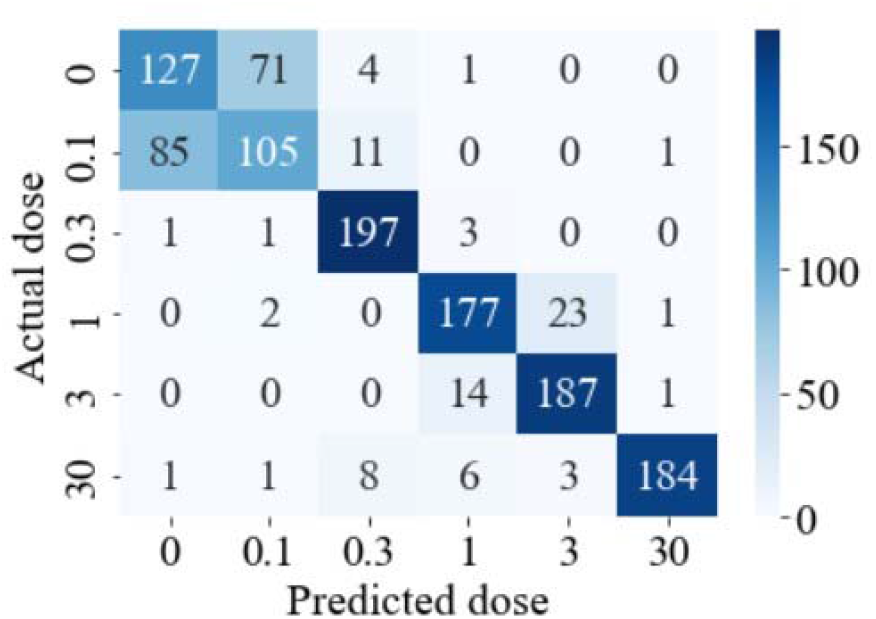
Confusion matrix of the deep neural network model using 5-channel image stacks as input for dose classification (0, 0.1, 0.3, 1, 3, 30 µM), ID validation.

**Table 1.**
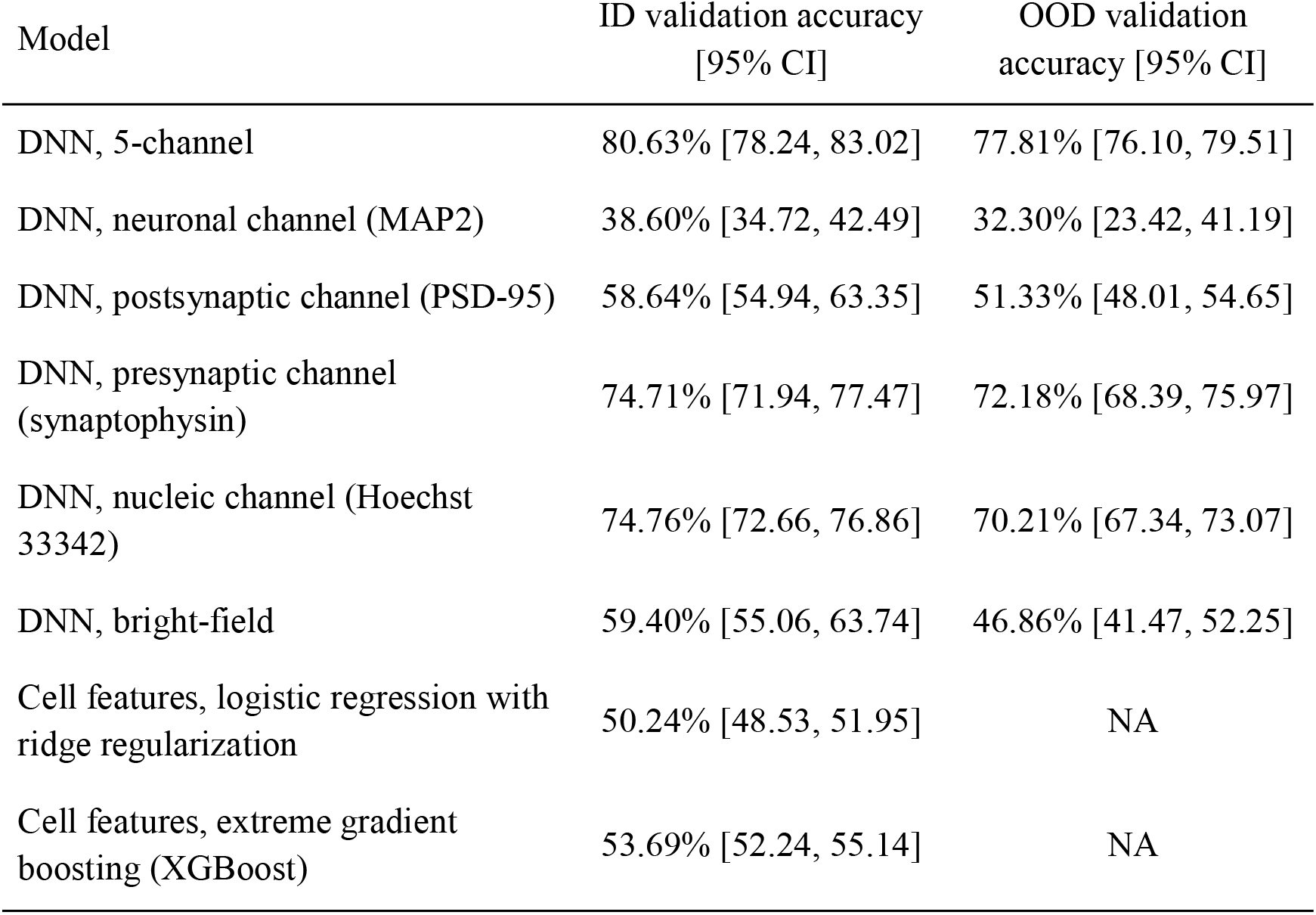
Model performance evaluation using accuracy (6 doses, i.e., 0, 0.1, 0.3, 1, 3, 30 µM). The DNN architecture is MobileNetV2 based. DNN: deep neural network. CI: confidence interval, calculated as the 2.5th to the 97.5th percentile of bootstrap estimates.

We have conducted several sensitivity analyses. Firstly, to further understand the contribution of each channel, we developed another 5 DNNs, each of which was trained and tested on images of one individual channel only. Results (Table 1) showed that the presynaptic marker (synaptophysin) and nuclear stain (Hoechst 33342) were the best performing single markers, both of which reached almost 75% accuracy, followed by the postsynaptic marker (PSD-95) at 58.64%. Somewhat surprisingly, the neuronal cytoskeleton marker (MAP2) yielded the worst accuracy of less than 40%. All models showed a marginal drop in accuracy in terms of out-of-distribution (OOD) validation, ranging from reasonable values of approximately 3% loss for the 5-channel, presynaptic and nucleic channel models, to roughly 6% loss for the rest. The model using the bright-field channel only achieved 59.40% on ID validation accuracy. However, the accuracy dropped significantly to 46.86% on OOD validation. We report the rest of the sensitivity analysis results in Table S3 in the supplement. Secondly, to confirm that the performance was entirely lost when the association between input images and their labels was eliminated, we conducted permutation training. During permutation training, the dose labels were shuffled and randomly assigned to image stacks. Not surprisingly, the accuracy of training with randomly shuffled labels dropped significantly to 16.41%, which was close to random guessing performance of 16.67%. Thirdly, to confirm that the DNN was not simply using information coming from the distribution of pixel intensity (mean, median, standard deviation, etc.) rather than the cellular features for decision making, we conducted pixel randomisation, where the pixels within each test image were randomly shuffled.

The accuracy of the pixel randomisation analysis was 16.52%, which was close to random guessing. Lastly, in addition to a well-established convolutional-based architecture (CNN), we tested the Transformer-based architecture and achieved comparable but slightly lower accuracy at 78.38%. We also evaluated the inference time of the trained CNN-based model and found that the network was able to process 305 images per second on a server with an Intel Xeon Silver 4216 CPU and a single GeForce RTX 2080 Ti graphical processing unit (GPU) accelerator.

### 2.2 Model interpretation

The saliency maps per channel are shown in Figure 3. According to saliency maps, the model was mainly focused on the area where the majority of cells were located. The results validated that the decision was made based on the actual cellular morphology, rather than being misled by the background irrelevant noise, which together with the previous OOD validation results indicates that there was no detectable batch effect between replicated wells across plates. Moreover, when pixels of each test image were randomly shuffled, the model’s accuracy dropped to 16.52%/16.40% in terms of ID and OOD validation (Table 2), indicating that the DNN was not simply measuring the statistics of pixel value intensities (mean, median, standard deviation, etc.) to perform classification. It is noteworthy that the model demonstrated the capacity to focus on the most pertinent areas of interest, rather than simply learning to identify cells or their general density. For instance, in Figure 3C, the model disregarded similar cells and concentrated solely on a specific region (the right area), which was closely linked to the cell phenotype resulting from the treatment. Moreover, to visualize the distribution of learned cellular representations, we employed the Uniform Manifold Approximation and Projection (UMAP) technique to transform the 512-dimensional feature vector into a 2-dimensional representation, depicted in Figure 4. The findings align well with the confusion matrix presented in Figure 2. For instance, the model primarily struggled to differentiate between 0 and 0.1 µM doses, as evidenced by the low accuracy in Figure 2 and the extensively overlapping points in Figure 4.

**Figure 3.**
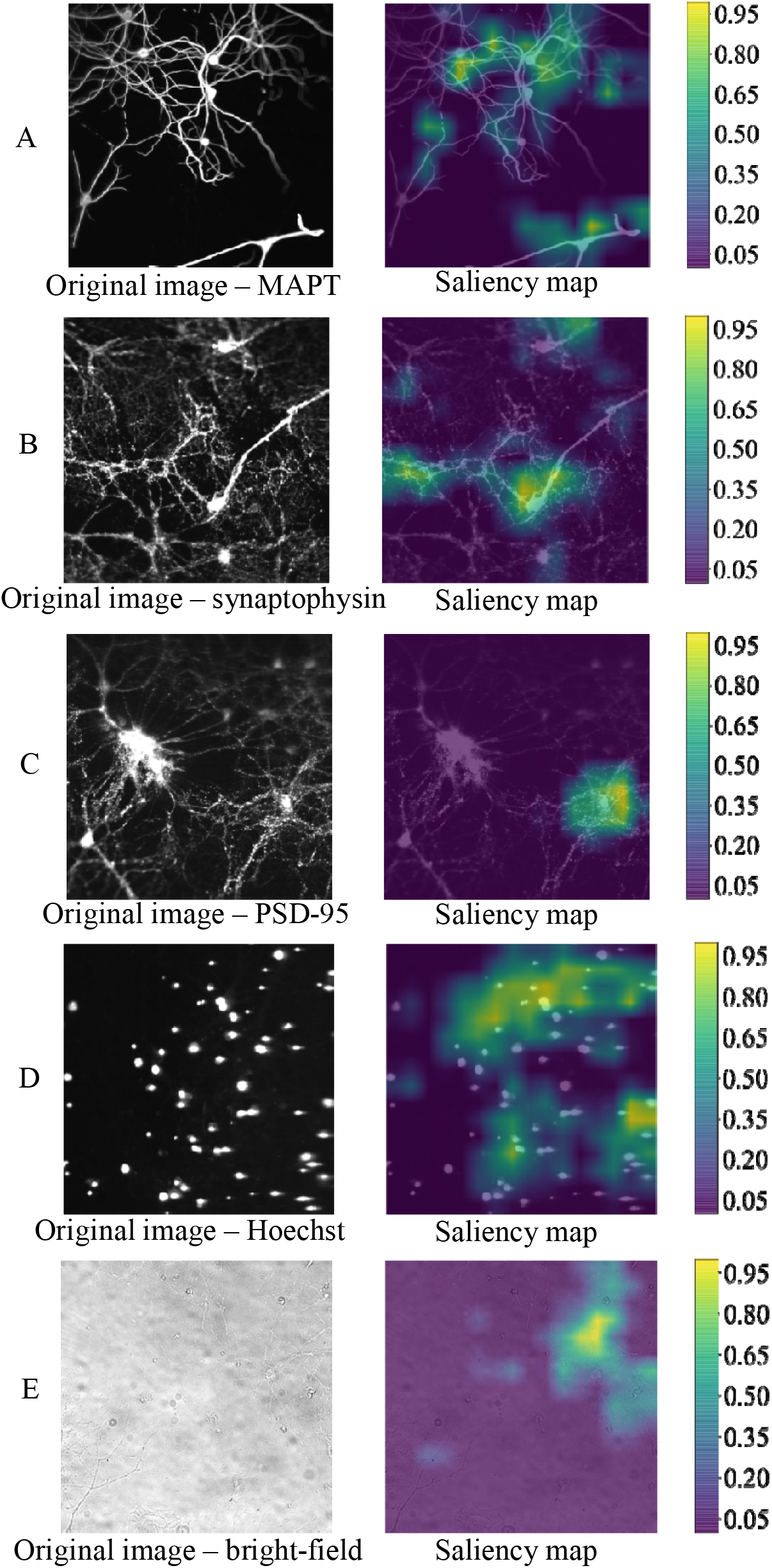
Grad-CAM images. A. MAPT immunostaining B. Synaptophysin immunostaining. C. PSD-95 immunostaining. D. Hoechst stain. E. Bright-field. Grad-CAM: Gradient-weighted Class Activation Mapping. A saliency map is an image that highlights the region on which the DNN focus in order to reflect the degree of importance of a pixel to the DNN model. According to the saliency maps, the model was mainly focused on the area where the majority of cells were distributed. The results validated that the decision was made based on the actual cellular morphology, rather than being misled by the background irrelevant noise.

**Figure 4.**
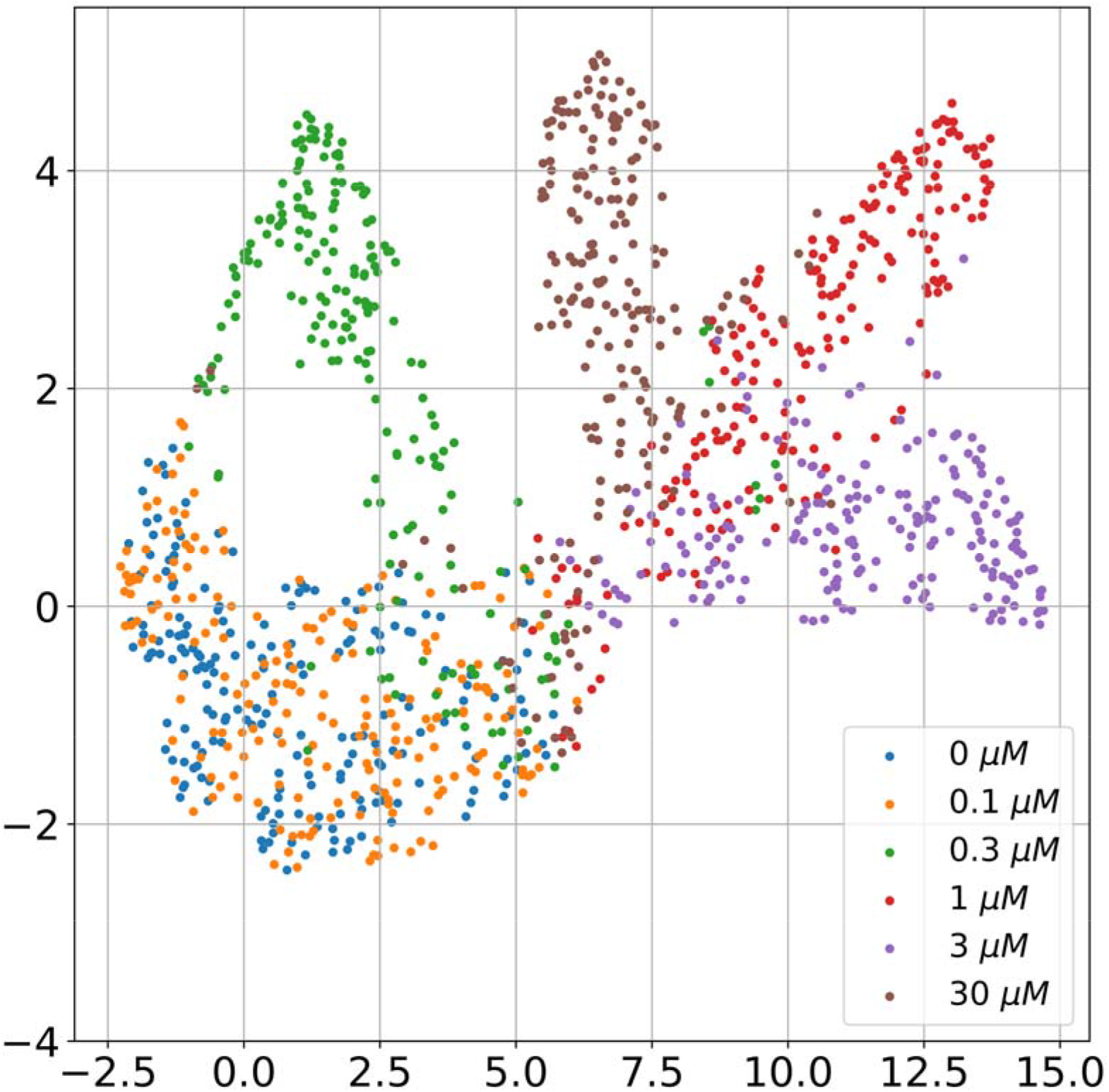
The UMAP projection of 1280-dimensional representation of cellular features. Some classes, denoted by doses in µM, demonstrate good separation, while others were challenging for the model to distinguish clearly. The results are well aligned with the confusion matrix shown in.

**Table 2.** Performance of our model and 2 biologists on the randomly selected 120 image stacks.

|  | Accuracy |
| --- | --- |
| Our model | 85.00% |
| Biologist 1 | 52.50% |
| Biologist 2 | 33.33% |

### 2.3 Cell feature analysis

We report the accuracy of logistic regression and XGBoost which used cell morphological features as input to classify Aβ_1-42_ doses in Table 1. These features were extracted by the Columbus software ^24^ from the same images as those used by the DNNs. Compared to logistic regression, the XGBoost achieved slightly higher accuracy at 53.69% in terms of ID validation, indicating that some non-linear relationships were found among the cell features. However, it still performed approximately 35% worse than the DNN (80.63%). The confusion matrices of these two models are shown in Figures S1 and S2 in the supplement. Misclassifications were mainly among neighbouring doses. Particularly, compared to the DNN, both models struggled to differentiate 0.1 and 0.3 µM doses from vehicle, as well as 30 µM from 1 and 3 µM.

We used SHAPley Additive exPlanations (SHAP) values to visualise feature importance^25 26^. A bar plot of the 20 most important features for the XGBoost model demonstrated by SHAP values is shown in Figure 5. The impact of each cell feature on the prediction of all 6 doses is stacked. The y-axis indicates the cell feature names sorted by importance. The x-axis is the stacked mean absolute value of the SHAP values of each feature. Overall, the standard deviation of ‘synaptic spot background intensity’ and ‘synaptic uncorrected spot peak intensity’, the mean of ‘postsynaptic total spot area’ and the max of ‘synaptic region intensity’ are the dominant cell morphological features. We can also see each feature contributed equally to negative control and the 0.1 µM Aβ_1-42_ (colours olive and violet) in most cases. This corresponds to the results in confusion matrices, i.e., misclassification between them is relatively high. The feature analyses for each specific dose can be seen in Figures S6 – S11 in the supplement, which reveals, for example, that a high standard deviation of the ‘synaptic spot background intensity’ lowers the predicted probability for the 0.1 µM Aβ_1-42_.

**Figure 5.**
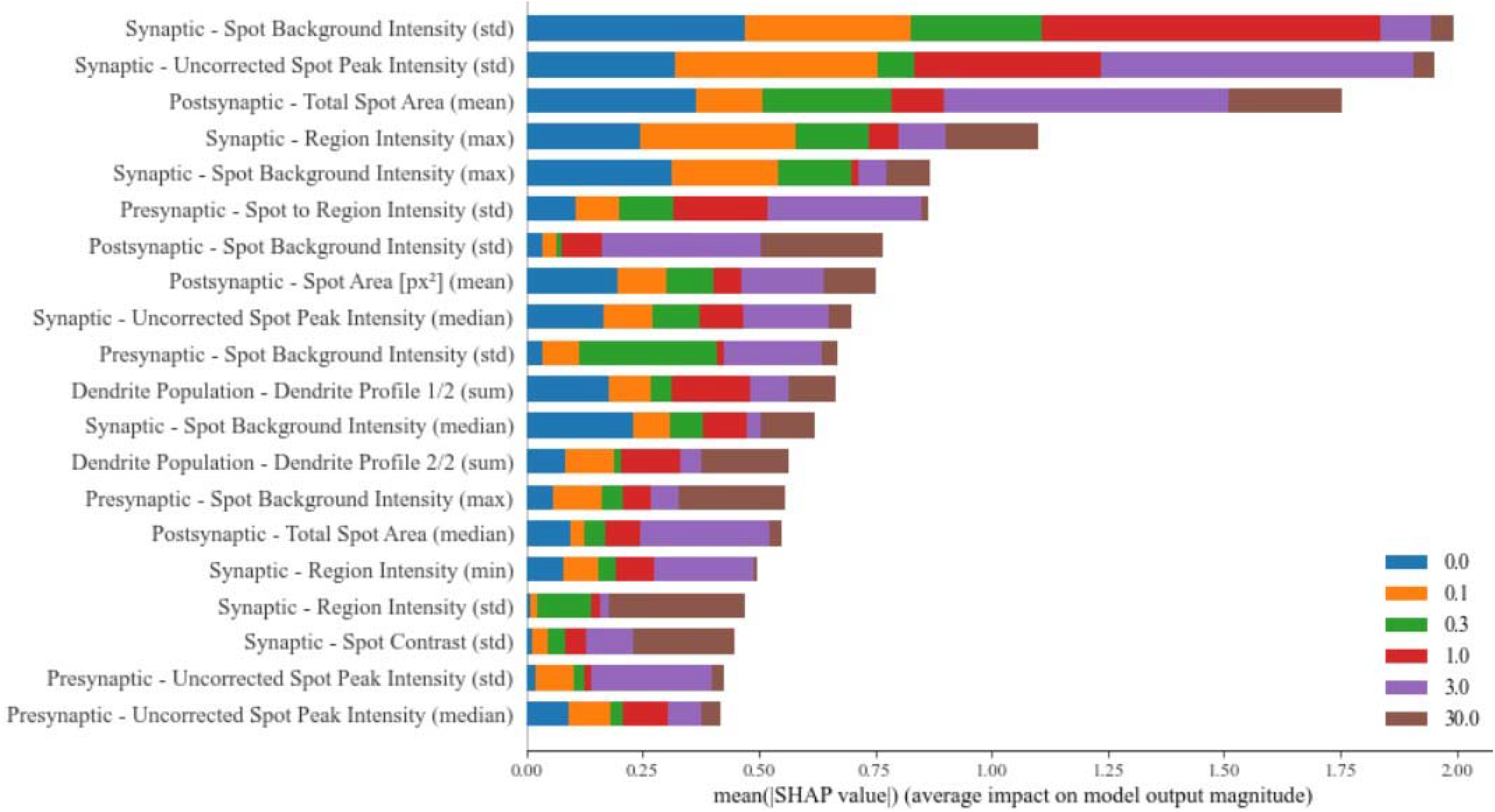
20 most important features for the XGBoost model measured by SHAP values. The impact of each cell feature on the prediction of all 6 doses is stacked to create the feature importance visualization. The y-axis indicates the cell feature names sorted by the importance. The x-axis is the stacked mean absolute value of the SHAP values of each feature.

### 2.4 Comparison of deep learning to human classification

We compared the performance of our model to this of 2 human experts on 120 image stacks that were randomly selected and manually annotated (see Table 2). Results show that our model performed better with 85% accuracy, while humans performed at 53% and 33%. Inter-annotator agreement had a Cohen’s kappa value of 0.34. According to the interpretation guidelines ^27^, this indicates a fair/poor agreement between human annotators, i.e., in this task humans made different annotation decisions for most doses and the task was hard. The confusion matrices are shown in Figures S3, S4 and S5 in the supplement. The majority of misclassifications occurred among adjacent doses.

## 3. Discussion

In this study, we present an end-to-end image-based cell profiling approach, i.e., a deep learning model that can reliably identify morphological changes in neuronal cells treated with a range of concentrations of Aβ_1-42_. The model outperformed traditional high content imaging tools and domain experts, while the OOD validation results proved the generalisability of the model. According to the DNNs trained using individual channels only, information carried by the presynaptic (synaptophysin) and nuclear markers (Hoechst 33342) was dominant for decision making. This was also confirmed by the results from the conventional HCI analysis (the XGBoost model) that, although achieving significantly lower accuracy, identified that the most important features were synapse related, particularly postsynaptic features marked by PSD-95. One of the earliest pathological events to occur in Alzheimer’s disease is synaptic dysregulation and loss ^28 29^. The results here suggest this remains a key feature, and consequently is a relevant model of disease able to utilise low doses of Aβ_1-42_.

Interestingly, the ability to classify using nuclear markers (Hoechst 33342) was not simply a result of focusing on the density of nucleus, as random zoom augmentation (images were randomly zoomed in or out by a threshold of 20% along both the height and width) was applied during training, which can also be seen from our OOD validation results. None of the models could distinguish between vehicle and 0.1 µM Aβ_1-42_. Other antibodies or imaging probes, recognising different cellular compartments may be necessary to discriminate among the subtle morphological changes produced by these adjacent doses of Aβ_1-42._ Such candidate agents may include markers of organelles that are implicated in Aβ_1-42_ metabolism, including ER/Golgi, endosomes, lysosomes ^30^.

We noticed a slight performance drop when we used the Transformer-based architecture. One possible reason is that unlike other tasks, cells in our images did not have a dominant pattern. Thus, splitting an image into patches and projecting these flattened patches yielded similar results across all patches, which diminish the gain of patch embeddings. Another reason might be that we had employed the same pre-training and fine-tuning hyperparameters for both CNNs and Transformers. Research has shown that CNNs can be at least as robust and expressive as the Transformers ^31^. Future work might include hand-crafting different learning rate scheduler or trying the hybrid CNN-Transformer architecture, but this is beyond the focus of this paper.

The work presented in this manuscript utilizes primary neurons isolated from C57BL/6J mice which is one of the most widely used inbred strains of mouse models and has been studied in various research areas. Primary murine neurons are a widely accepted model employed in genetics and neuroscience research, as a proof-of-concept to demonstrate the power of AI/ML-enabled models in exploring cellular pathobiology. The model developed in this work can serve as a foundation to test the transferability to human cellular systems (e.g., hiPSC-derived neuronal models) carrying various genotypes and/or with pathological insults. These cellular models may be more relevant and have better translatability to human compared to rodent models that have been traditionally used in preclinical research.

In conclusion, we have shown that end-to-end deep learning models can identify Aβ_1-42_ treatment conditions based on the morphological changes of neuronal cells. The speed and accuracy of our deep learning model outperformed conventional cell profiling methods and human experts and successfully classified changes in phenotype in response to a range of Aβ_1-42_ concentrations. More generally, in this work, using a dose-response case study, we generated a range of cell phenotypes that are close and yet distinguishable by deep learning model, and demonstrated the potential of machine learning as a valuable tool to recognise changes in cell phenotypes induced in response to drug treatment. The ability to robustly learn and rapidly recognise phenotypic changes in cells can be utilised in large scale screening procedures with chemogenomic libraries and identify those compounds that block or reverse the neurodegenerative phenotype, thereby accelerating the discovery of novel therapeutic targets.

## 4. Methods

### 4.1 Data acquisition

Cortical neurons were prepared from E15 embryos harvested from C57BL/6J mice in accordance with the Animals Scientific Procedures Act (1986). Pregnant mice were killed by CO_2_, followed by cervical dislocation. Embryos were removed and killed by cervical dislocation, and cortices were dissected into Trypsin-EDTA. After washing and treating with DNase, cortices were triturated to a single cell suspension, centrifuged and resuspended in Neurobasal medium with MACs Neurobrew supplement and L-Glutamine. Cells were then plated in poly-D-lysine precoated 96 well plates at a density of 50,000/cm^2^.

Mouse primary neuronal cultures were then treated with Aβ_1-42_ in 6 doses (0, 0.1, 0.3, 1, 3, 30 µM) and immunostained with Hoechst 33342 and antibodies that recognised microtubule-associated protein 2 (MAP2), synaptophysin and postsynaptic density-95 (PSD-95) to allow visualisation of the nucleus, neuronal cytoskeleton and localisation of pre- and post-synaptic markers. These markers were selected to enrich information related to both general cell morphology and to neuronal functionality, both of which modalities are known to be altered in response to β-amyloid. The experiment was replicated on 5 plates, with 54 wells per plate (9 replications per Aβ_1-42_ dose). The cells were finally imaged at 40x magnification and 225 different field-of-views per well were captured. We obtained 60,750 images at 12,150 different locations on each plate. At each location, four immunostained images and one bright-field image were captured. All images were grayscale with a resolution of 1024 x 1024.

### 4.2 Model design

The main models were deep neural networks (DNN), i.e., a deep learning-based machine learning model inspired by our brain’s network of neurons, that can identify morphological changes of neuronal cells treated with 6 concentrations of Aβ_1-42_. A set of additional DNNs were developed for further performance evaluation and model interpretation, i.e., identifying the features in stained images that contributed the most to the model for decision making. The workflow is shown in Figure 1.

Subsequent to image capture, all fluorescent and bright-field greyscale images from each unique field of view were used to produce a stack of 5-channel image, thereby forming the input data structure of the model. Pixel intensity values were then normalised. In terms of the training pipeline (represented by blue arrows in Figure 1), image augmentation techniques such as random flip, rotation and zoom, were applied to create transformed versions of images to artificially expand the dataset, which enabled our model to be more robust by introducing potential extra variations in the real world and prevent model overfitting. The image augmentation was applied only on the training dataset rather than the validation or testing sets, and the augmentation was re-calculated for each batch at the beginning of each epoch. We deployed the MobileNetV2 (53 layers, 3.4 million parameters) for feature extraction, which is a lightweight DNN architecture with fewer parameters and is designed for mobile platforms ^32^. The original input layer of MobileNetV2 (3 channels only) was replaced by our data structure (1024 x 1024 x 5, i.e., 5 channels). The original classification layer was removed, and a max pooling layer and global average pooling layer were attached afterwards. Transfer learning was applied using parameters pre-trained on ImageNet ^33^. The other layers were initialised using Glorot’s initialisation ^34^. TensorFlow was used for implementation ^35^. We used 80% images for training, 10% for validation and the rest 10% for testing. Fine-tuning was carried out in two steps. In the first step, we fixed the parameters of MobileNetV2 and only updated the parameters of the other layers using 30 epochs, 16 mini-batch size and 0.0002 learning rate. We then unfroze the MobileNetV2 and fine-tuned all layers using 0.00001 learning rate and 10 early stopping patience. All other hyperparameters were reported in Table S1 in the supplement. The model adopted an end-to-end training manner, where it took a raw 5-channel image stack and directly yielded 6 numbers, each corresponding to the probability of treatment with the corresponding dose. Therefore, the task was essentially a supervised classification problem.

Gradient-weighted Class Activation Mapping (Grad-CAM) was deployed for model interpretation, where the gradient flowing into the last convolutional layer and the feature maps produced by the last convolutional layer were combined to generate a heatmap indicating the focus of our model when making decisions ^36^. Grad-CAM typically results in a saliency map with a lower resolution than the input image. Thus, we followed the paper and used bilinear interpolation to generate the saliency maps ^36^. In this study, single-channel images were fed into the DNN after training was completed and the parameters were fixed to obtain Grad-CAM images (orange arrows in Figure 1). We then overlay them on original images for clearer visualisation.

To further evaluate the main model, we performed four sensitivity analyses. Firstly, to further understand the contribution of each channel, we developed another 5 DNNs, each of which was trained and tested on images of one individual channel only (4 stained and bright-field). The architecture and hyperparameters of these 5 DNNs were the same as in their 5-channel equivalent described above, with the only difference that the input layer had only 1 channel. Secondly, we performed permutation training, where the dose labels were shuffled and randomly assigned to image stacks, to confirm that all performance was lost when all association between input images and their labels was eliminated. Thirdly, we conducted pixel randomisation, where the pixels within each test image were randomly shuffled, to confirm that the DNN was not simply using information coming from the distribution of pixel intensity (mean, median, standard deviation, etc.) rather than the cellular features for decision making. Lastly, instead of fine-tuning MobileNetV2 which is a convolutional-based architecture, we tested another Transformer-based architecture – DeiT-Ti with similar number of parameters (5 million parameters, 3 heads, 12 layers, 192 embedding dimensions, 16 patches) ^37^. Other hyperparameters were the same as the MobileNetV2-based DNN, except we used cosine learning rate decay in the fine-tuning phase. Detailed hyperparameters are in Table S1 in the supplement.

### 4.3 Cell feature analysis

To compare the end-to-end DNN model with traditional cell profiling methods ^17^, we built two conventional models and used cell morphological features as input to classify the doses. Specifically, using the channels allocated for each marker Hoechst, Synaptophysin, PSD-95 and MAP2 a customised pipeline was developed on the Columbus Image Data Storage and Analysis System ^24^ and used to determine 302 morphological features focusing on changes in nuclear, neurite and synaptic number, expression intensity, and morphology.

Nuclear morphological features were determined by the corresponding channel using ‘Find Nuclei’ on Method B, removing border objects. Standard intensity properties, standard and STAR morphology properties were then calculated for each object. Features relating to neurites were obtained by identifying MAP2-positive neurites as the region of interest population using a common threshold of 0.5. The area, standard intensity properties, standard and STAR morphology properties were calculated for each field. To analyse synaptic changes, The MAP2 region was resized by 7px and using Method A of Find Spots to identify the pre-synaptic and post-synaptic channels along the MAP2 region. Signal from both pre- and post-synaptic channels was excluded if over 100px^2^. To identify synapses, the overlap of the pre- and post-synaptic spots was determined by resizing the outer border of the pre-synaptic spots. The number of objects, and features pertaining to spot intensity were determined for pre-, post-synaptic, and combined channels.

For each characteristic identified per objects per field of view (same as the field of views used for the DNN model), we quantified each of the neurite, synaptic and nuclei related characteristics by 6 statistical metrics, namely sum, mean, standard deviation, median, minimum and maximum values. This excluded dendrite related characteristics, which were quantified by sum only due to limited capacity to accurately segment individual neurons. A detailed list of each characteristic identified using Columbus software can be found in Table S2 in the supplement.

We then built two models, namely a statistical model (logistic regression with ridge regularisation) and a machine learning model (extreme gradient boosting, XGBoost) ^38^, and used these features as model input for classification. All features were standardized, i.e., rescaled to have a mean of 0 and a standard deviation of 1. We employed distributed asynchronous hyper-parameter optimisation (Hyperopt) for hyper-parameter tuning, i.e., choosing a set of optimal arguments for a model whose values are set before the training process. Finally, we performed model interpretation for the XGBoost by using SHAP values to demonstrate the global feature importance and the impact of each feature on individual dose specific predictions ^25 26^. SHAP value is the difference between predicted probability and base value, given a list of features. The base value is the value that will be predicted if we do not know any features for an image stack, i.e., the average of the model output over the training dataset. A higher magnitude of a SHAP value indicates a more important and predictive feature.

### 4.4 Comparison of deep learning and human classification

We further compared our model (6 doses classification using 5-channel image stacks) to human experts. In this study, two biologists, each experienced in microscopic analysis of Aβ_1-42_ treated neurons, were invited to carry out annotation, i.e., the process of manually labelling images. Specifically, the biologists were first given sufficient images to become familiar with the task (training). In our case, 120 image stacks, 20 per dose, were provided. Each of the experts then received 120 image stacks, 20 per dose, with random image names and was tasked with predicting the dose of each image stack. The testing images provided to both experts were identical. We finally compared the performance of the biologists to our model. Accuracy was used for evaluation, which is detailed in the subsequent section. In addition, we calculated inter-annotator agreement, which is a measure of how well multiple annotators can make the same decision for a specific category. In our study, this indicates how easy it is to clearly distinguish the doses based on cell morphological changes by human experts. Cohen’s kappa statistic was used to measure the inter-annotator agreement ^39^.

### 4.5 Model evaluation setting

Both ID and OOD validations were performed. In our case, we considered using images from the same plate of training images for testing as ID validation, whereas OOD validation was defined by training the DNN using images from one plate and testing on another plate.

In terms of model evaluation, bootstrapping (a statistical procedure that resamples a single dataset to create many simulated samples by random sampling with replacement) was deployed, where images, image stacks or cell features from one plate were randomly drawn with replacement to form a training set. The trained model was then tested on the testing set, i.e., images of the same plate for ID validation. This process was repeated 20 times with random initialisation each time. All DNNs were validated on OOD images randomly drawn from another plate, with the total number of test images identical to the ones used for ID validation. The same training and evaluation strategies were applied for the cell feature analysis with logistic regression and XGBoost. No OOD validation was performed for the cell feature analysis.

Accuracy was used to evaluate all models and was calculated by:

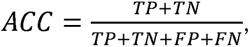

where TP, TN, FP and FN represent true positive, true negative, false positive and false negative, respectively. As a by-product, confusion matrices were generated as well for stratified misclassification identification.

All DNNs were developed and validated using TensorFlow ^35^. Logistic regression, XGBoost and SHAP were implemented in Python ^40^.

## Data Availability

The data and codes that support the findings of this study are available on request from the first and corresponding authors.

## Acknowledgements

NJB, ANH and AK would like to acknowledge the support from GSK Functional Genomics research grant. A.K. were supported in part by the NIHR AI Award for Health and Social Care (AI_AWARD02183). The views expressed are those of the authors and not necessarily those of the University of Oxford, the University of Bristol or GlaxoSmithKline.

## Author contributions

QL and HR were responsible for the analysis. FN, HR, and ER were responsible for data acquisition. HR and ADV carried out annotation. QL, HR, ANH, NJB and AK were responsible for study conception. HR, SL and NJB provided biological interpretation. ANH, NJB and AK supervised the project. All authors contributed to the manuscript.

## References

1. Tomita M. Whole-cell simulation: a grand challenge of the 21st century. Trends in Biotechnology 2001;19(6):205–10.

2. Carragher N, Piccinini F, Tesei A, et al. Concerns, challenges and promises of high-content analysis of 3D cellular models. Nature Reviews Drug Discovery 2018;17(8):606–06.

3. Avior Y, Sagi I, Benvenisty N. Pluripotent stem cells in disease modelling and drug discovery. Nature Reviews Molecular Cell Biology 2016;17(3):170–82.

4. Gitler AD, Dhillon P, Shorter J. Neurodegenerative disease: models, mechanisms, and a new hope. Disease Models & Mechanisms 2017;10(5):499–502.

5. Lin Y-T, Seo J, Gao F, et al. APOE4 causes widespread molecular and cellular alterations associated with Alzheimer’s disease phenotypes in human iPSC-derived brain cell types. Neuron 2018;98(6):1141–54. e7.

6. Fernando MB, Ahfeldt T, Brennand KJ. Modeling the complex genetic architectures of brain disease. Nature Genetics 2020;52(4):363–69.

7. Wiegand C, Banerjee I. Recent advances in the applications of iPSC technology. Current Opinion in Biotechnology 2019;60:250–58.

8. Dementia - World Health Organization March 2021 [Available from: https://www.who.int/news-room/fact-sheets/detail/dementia accessed 17 August 2021.

9. Canter RG, Penney J, Tsai L-H. The road to restoring neural circuits for the treatment of Alzheimer’s disease. Nature 2016;539(7628):187–96.

10. Colom-Cadena M, Spires-Jones T, Zetterberg H, et al. The clinical promise of biomarkers of synapse damage or loss in Alzheimer’s disease. Alzheimer’s Research & Therapy 2020;12(1):1–12.

11. Murphy MP, LeVine III H. Alzheimer’s disease and the amyloid-β peptide. Journal of Alzheimer’s Disease 2010;19(1):311–23.

12. Hyman BT, Phelps CH, Beach TG, et al. National Institute on Aging–Alzheimer’s Association guidelines for the neuropathologic assessment of Alzheimer’s disease. Alzheimer’s & Dementia 2012;8(1):1–13.

13. Hammond TC, Xing X, Wang C, et al. β-amyloid and tau drive early Alzheimer’s disease decline while glucose hypometabolism drives late decline. Communications Biology 2020;3(1):1–13.

14. Lee J-H, Yang D-S, Goulbourne CN, et al. Faulty autolysosome acidification in Alzheimer’s disease mouse models induces autophagic build-up of Aβ in neurons, yielding senile plaques. Nature Neuroscience 2022:1–14.

15. Kaur G, Dufour JM. Cell lines: Valuable tools or useless artifacts: Taylor & Francis, 2012:1–5.

16. Iqbal MS, Ahmad I, Bin L, et al. Deep learning recognition of diseased and normal cell representation. Transactions on Emerging Telecommunications Technologies 2021;32(7):e4017.

17. Caicedo JC, Cooper S, Heigwer F, et al. Data-analysis strategies for image-based cell profiling. Nature Methods 2017;14(9):849–63.

18. Mzurikwao D, Khan MU, Samuel OW, et al. Towards image-based cancer cell lines authentication using deep neural networks. Scientific Reports 2020;10(1):1–15.

19. Gibson LC, Shin BC, Dai Y, et al. Early leptin intervention reverses perturbed energy balance regulating hypothalamic neuropeptides in the pre-and postnatal calorie-restricted female rat offspring. Journal of Neuroscience Research 2015;93(6):902–12.

20. Wainberg M, Merico D, Delong A, et al. Deep learning in biomedicine. Nature Biotechnology 2018;36(9):829–38.

21. Christiansen EM, Yang SJ, Ando DM, et al. In silico labeling: predicting fluorescent labels in unlabeled images. Cell 2018;173(3):792–803. e19.

22. Al-Kofahi Y, Zaltsman A, Graves R, et al. A deep learning-based algorithm for 2-D cell segmentation in microscopy images. BMC Bioinformatics 2018;19(1):1–11.

23. Falk T, Mai D, Bensch R, et al. U-Net: deep learning for cell counting, detection, and morphometry. Nature Methods 2019;16(1):67–70.

24. PerkinElmer. Columbus Image Data Storage and Analysis System [Available from: https://www.perkinelmer.com/uk/product/image-data-storage-and-analysis-system-columbus accessed 17 December 2021.

25. A unified approach to interpreting model predictions. Proceedings of the 31st International Conference on Neural Information Processing Systems; 2017.

26. Xie YR, Castro DC, Bell SE, et al. Single-cell classification using mass spectrometry through interpretable machine learning. Analytical Chemistry 2020;92(13):9338–47.

27. Viera AJ, Garrett JM. Understanding interobserver agreement: the kappa statistic. Family Medicine 2005;37(5):360–63.

28. Davies C, Mann D, Sumpter P, et al. A quantitative morphometric analysis of the neuronal and synaptic content of the frontal and temporal cortex in patients with Alzheimer’s disease. Journal of the neurological sciences 1987;78(2):151–64.

29. Terry RD, Masliah E, Salmon DP, et al. Physical basis of cognitive alterations in Alzheimer’s disease: synapse loss is the major correlate of cognitive impairment. Annals of Neurology: Official Journal of the American Neurological Association and the Child Neurology Society 1991;30(4):572–80.

30. Saido T, Leissring MA. Proteolytic degradation of amyloid β-protein. Cold Spring Harbor Perspectives in Medicine 2012;2(6):a006379.

31. An impartial take to the cnn vs transformer robustness contest. Computer Vision–ECCV 2022: 17th European Conference, Tel Aviv, Israel, October 23–27, 2022, Proceedings, Part XIII; 2022. Springer.

32. MobileNetV2: Inverted residuals and linear bottlenecks. Proceedings of the IEEE Conference on Computer Vision and Pattern Recognition; 2018.

33. Krizhevsky A, Sutskever I, Hinton GE. ImageNet classification with deep convolutional neural networks. Advances in Neural Information Processing Systems 2012;25:1097–105.

34. Understanding the difficulty of training deep feedforward neural networks. Proceedings of the thirteenth international conference on artificial intelligence and statistics; 2010. JMLR Workshop and Conference Proceedings.

35 TensorFlow: A system for large-scale machine learning. 12th USENIX Symposium on Operating Systems Design and Implementation; 2016.

36. Grad-CAM: Visual explanations from deep networks via gradient-based localization. Proceedings of the IEEE International Conference on Computer Vision; 2017.

37. Training data-efficient image transformers & distillation through attention. International Conference on Machine Learning; 2021. PMLR.

38. XGBoost: A scalable tree boosting system. Proceedings of the 22nd ACM SIGKDD International Conference on Knowledge Discovery and Data Mining; 2016.

39. Artstein R. Inter-annotator agreement. Handbook of linguistic annotation: Springer 2017:297–313.

40. Raschka S. Python machine learning: Packt Publishing Ltd 2015.

