## Supplemental material for "Deep learning-based cell profiling based on neuronal morphology"

### Model Details (Table S1)

Table S1. Implementation details of the deep neural network using 5-channel image stacks for 6 dose (0, 0.1, 0.3, 1, 3, 30 µM) classification.

| Structure | Image augmentation + MobileNetV2 + max pooling + global average pooling |
| --- | --- |
| Hyperparameter | Random flip: horizontal_and_vertical  Random rotation: 0.2  Random zoom: -0.2, 0.2  max pooling stride: 2  global average pooling dropout: 0.2  Initial epoch: 30  Learning rate: initial - 0.0002, fine-tuning - 0.00001  Optimiser: initial – Adam, fine-tuning - RMSprop  Loss: cross-entropy loss between the labels and predictions  Early stopping monitor: accuracy  Early stopping patience: 10 |

### Cell Features (Table S2)

Table S2. Cell features extracted through the Columbus Image Data Storage and Analysis System.

Synaptic - Relative Spot Intensity

Synaptic - Corrected Spot Intensity

Synaptic - Uncorrected Spot Peak Intensity

Synaptic - Spot Contrast

Synaptic - Spot Background Intensity

Synaptic - Spot Area [px²]

Synaptic - Region Intensity

Synaptic - Spot to Region Intensity

Synaptic - Number of Spots

Presynaptic - Relative Spot Intensity

Presynaptic - Corrected Spot Intensity

Presynaptic - Uncorrected Spot Peak Intensity

Presynaptic - Spot Contrast

Presynaptic - Spot Background Intensity

Presynaptic - Spot Area [px²]

Presynaptic - Region Intensity

Presynaptic - Spot to Region Intensity

Presynaptic - Number of Spots

Postsynaptic - Relative Spot Intensity

Postsynaptic - Corrected Spot Intensity

Postsynaptic - Uncorrected Spot Peak Intensity

Postsynaptic - Spot Contrast

Postsynaptic - Spot Background Intensity

Postsynaptic - Spot Area [px²]

Postsynaptic - Region Intensity

Postsynaptic - Spot to Region Intensity

Postsynaptic - Total Spot Area

Postsynaptic - Number of Spots

Dendrite Population - Dendrite Area [µm²]

Dendrite Population - Dendrite Roundness

Dendrite Population - Dendrite Width [µm]

Dendrite Population - Dendrite Length [µm]

Dendrite Population - Dendrite Ratio Width to Length

Dendrite Population - Dendrite Symmetry 02

Dendrite Population - Dendrite Symmetry 03

Dendrite Population - Dendrite Symmetry 04

Dendrite Population - Dendrite Symmetry 05

Dendrite Population - Dendrite Symmetry 12

Dendrite Population - Dendrite Symmetry 13

Dendrite Population - Dendrite Symmetry 14

Dendrite Population - Dendrite Symmetry 15

Dendrite Population - Dendrite Threshold Compactness 30%

Dendrite Population - Dendrite Threshold Compactness 40%

Dendrite Population - Dendrite Threshold Compactness 50%

Dendrite Population - Dendrite Threshold Compactness 60%

Dendrite Population - Dendrite Axial Small Length

Dendrite Population - Dendrite Axial Length Ratio

Dendrite Population - Dendrite Radial Mean

Dendrite Population - Dendrite Radial Relative Deviation

Dendrite Population - Dendrite Profile 1/2

Dendrite Population - Dendrite Profile 2/2

Nuclei - Nucleus Area [µm²]

Nuclei - Nucleus Roundness

Nuclei - Intensity Nucleus

Nuclei - Nucleus Symmetry 02

Nuclei - Nucleus Symmetry 03

Nuclei - Nucleus Symmetry 04

Nuclei - Nucleus Symmetry 05

Nuclei - Nucleus Symmetry 12

Nuclei - Nucleus Symmetry 13

Nuclei - Nucleus Symmetry 14

Nuclei - Nucleus Symmetry 15

Nuclei - Nucleus Threshold Compactness 30%

Nuclei - Nucleus Threshold Compactness 40%

Nuclei - Nucleus Threshold Compactness 50%

Nuclei - Nucleus Threshold Compactness 60%

Nuclei - Nucleus Axial Small Length

Nuclei - Nucleus Axial Length Ratio

Nuclei - Nucleus Radial Mean

Nuclei - Nucleus Radial Relative Deviation

Nuclei - Nucleus Profile 1/2

Nuclei - Nucleus Profile 2/2

### Model Performance Evaluation for the Sensitivity Analyses (Table S3)

Table S3. Model performance evaluation using accuracy. DNN: deep neural network. CI: confidence interval, calculated as the 2.5th to the 97.5th percentile of bootstrap estimates.

| Model | ID validation accuracy [95% CI] | OOD validation accuracy [95% CI] |
| --- | --- | --- |
| DNN, MobileNetV2 based, 5-channel, dose labels were randomly shuffled | 16.41% [14.97, 17.84] | 18.11% [14.76, 21.45] |
| DNN, MobileNetV2 based, 5-channel, pixels in test images were randomly shuffled | 16.52% [15.04, 17.99] | 16.40% [14.28, 18.51] |
| DNN, DeiT-Ti based, 5-channel | 78.38%, [74.90, 81.87] | 69.88%, [66.59, 73.17] |

### Confusion Matrices – Cell Feature Analysis (Figure S1, Figure S2)

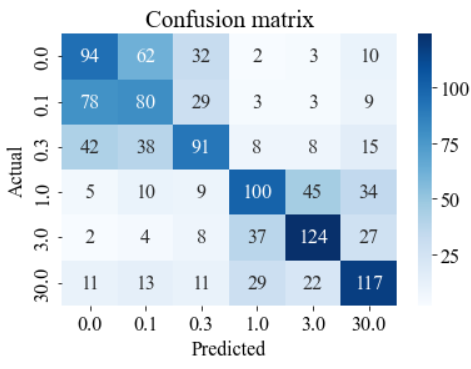

Figure S1. Confusion matrix of the logistic regression with ridge regularization using cell features as input for 6 dose (0, 0.1, 0.3, 1, 3, 30 µM) classification, internal validation.

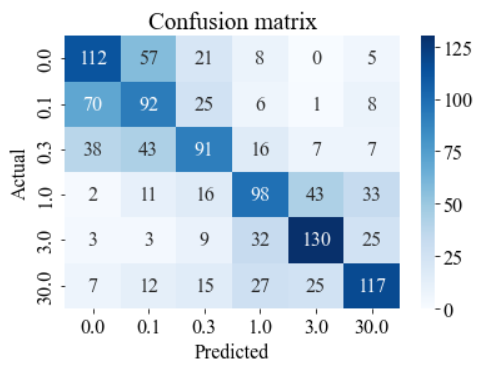

Figure S2. Confusion matrix of the extreme gradient boosting (XGBoost) using cell features as input for 6 dose (0, 0.1, 0.3, 1, 3, 30 µM) classification, internal validation.

### Confusion Matrices - Annotation (Figure S3, Figure S4, Figure S5)

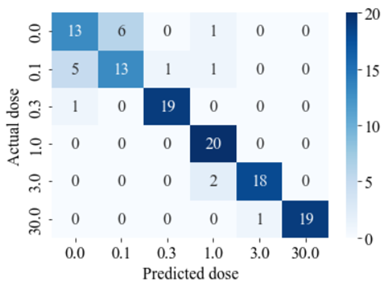

Figure S3. Confusion matrix of our model classifying annotation image stacks.

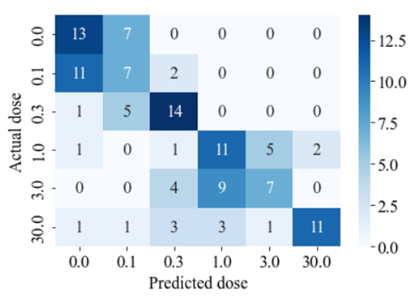

Figure S4. Confusion matrix of annotator 1 classifying annotation image stacks.

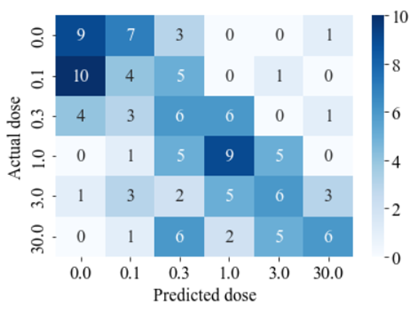

Figure S5. Confusion matrix of annotator 2 classifying annotation image stacks.

### Feature Importance Analysis of the XGBoost Model for Each Dose (Figure S6, Figure S7, Figure S8, Figure S9, Figure S10, Figure S11)

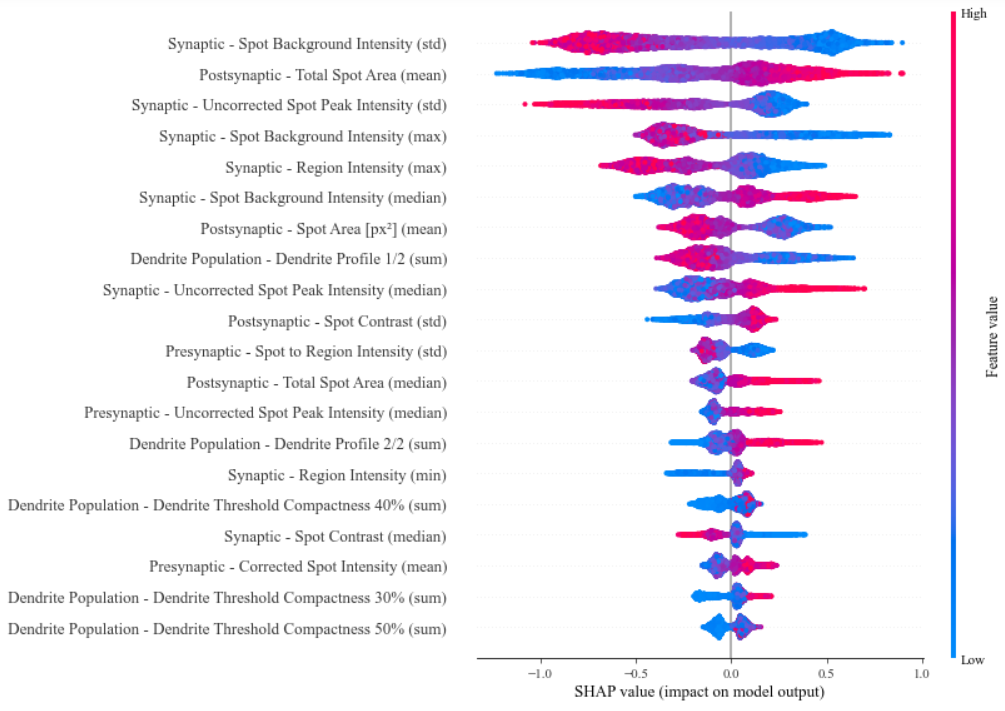

Figure S6. 20 most important features for the XGBoost model measured by SHAP values, dose 0. The y-axis indicates the cell feature names sorted by the importance. The x-axis is the SHAP values of each feature. The color represents the feature value (red high, blue low).

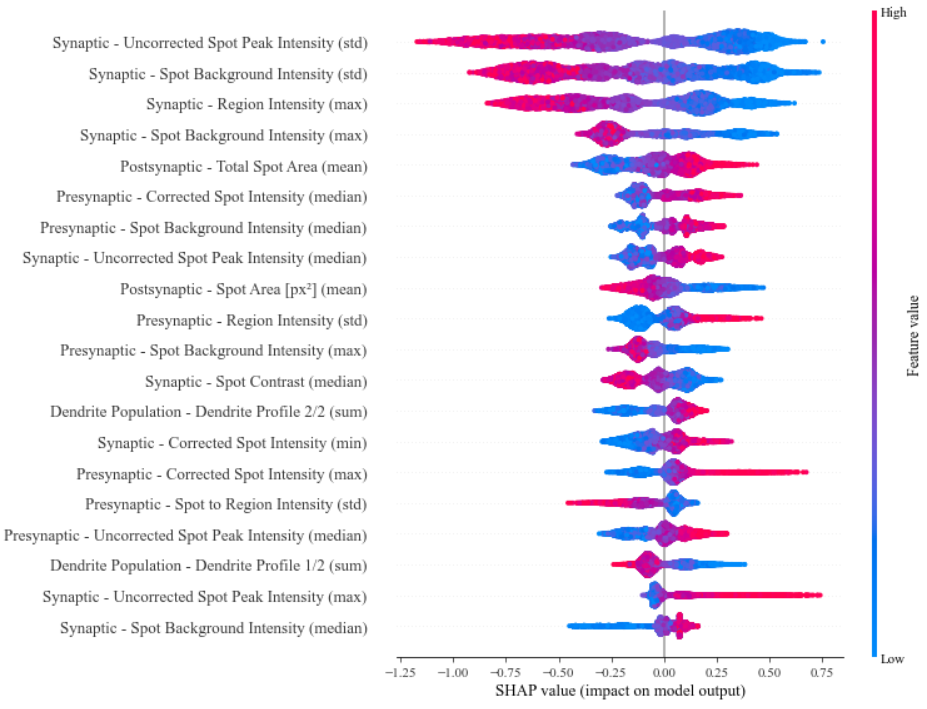

Figure S7. 20 most important features for the XGBoost model measured by SHAP values, dose 0.1. The y-axis indicates the cell feature names sorted by the importance. The x-axis is the SHAP values of each feature. The color represents the feature value (red high, blue low).

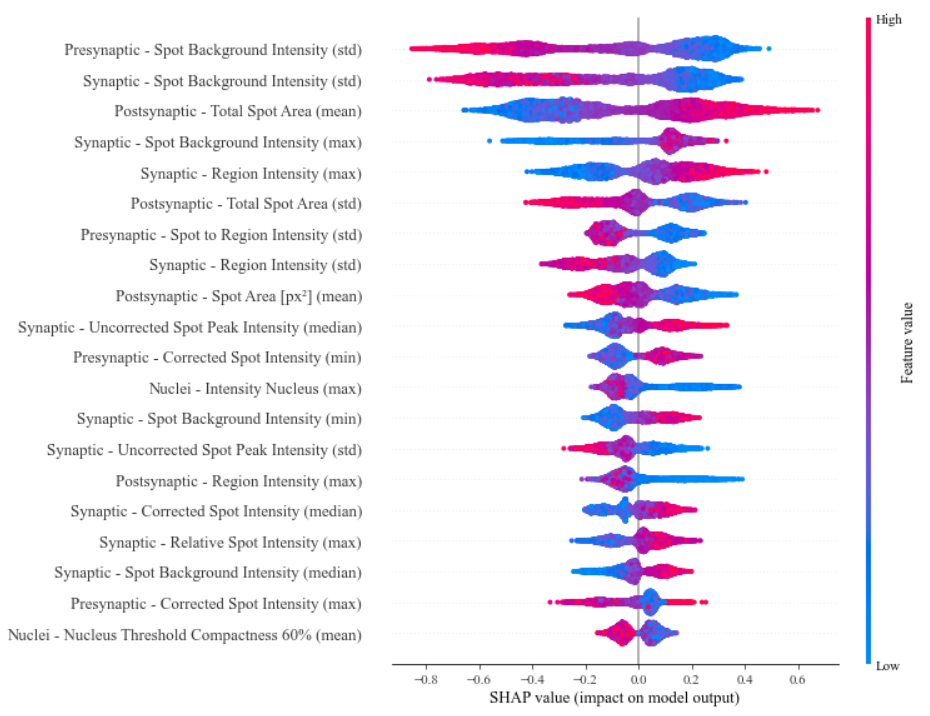

Figure S8. 20 most important features for the XGBoost model measured by SHAP values, dose 0.3. The y-axis indicates the cell feature names sorted by the importance. The x-axis is the SHAP values of each feature. The color represents the feature value (red high, blue low).

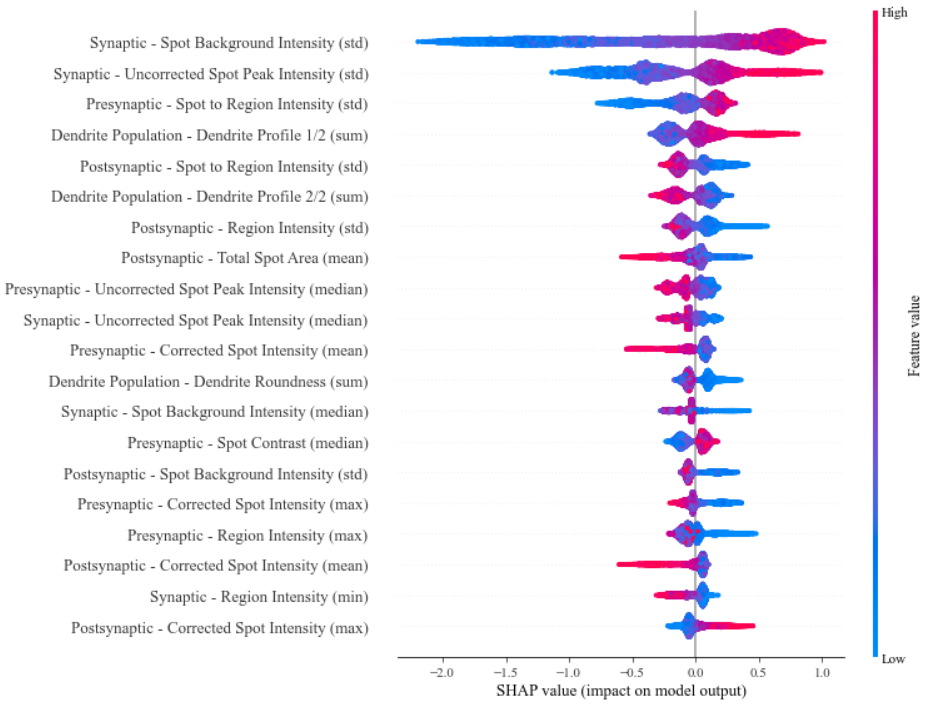

Figure S9. 20 most important features for the XGBoost model measured by SHAP values, dose 1. The y-axis indicates the cell feature names sorted by the importance. The x-axis is the SHAP values of each feature. The color represents the feature value (red high, blue low).

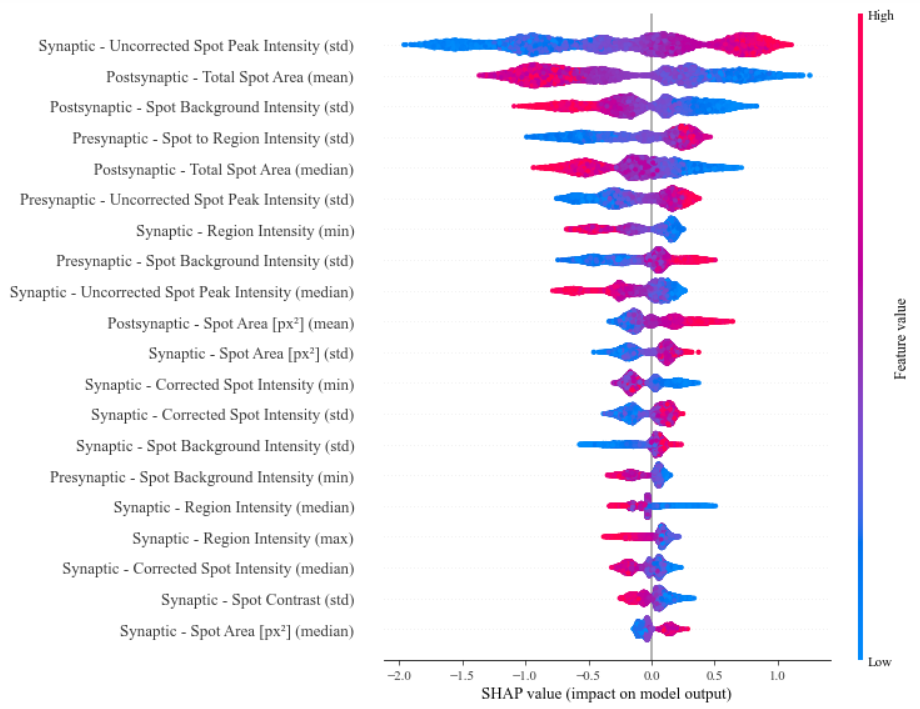

Figure S10. 20 most important features for the XGBoost model measured by SHAP values, dose 3. The y-axis indicates the cell feature names sorted by the importance. The x-axis is the SHAP values of each feature. The color represents the feature value (red high, blue low).

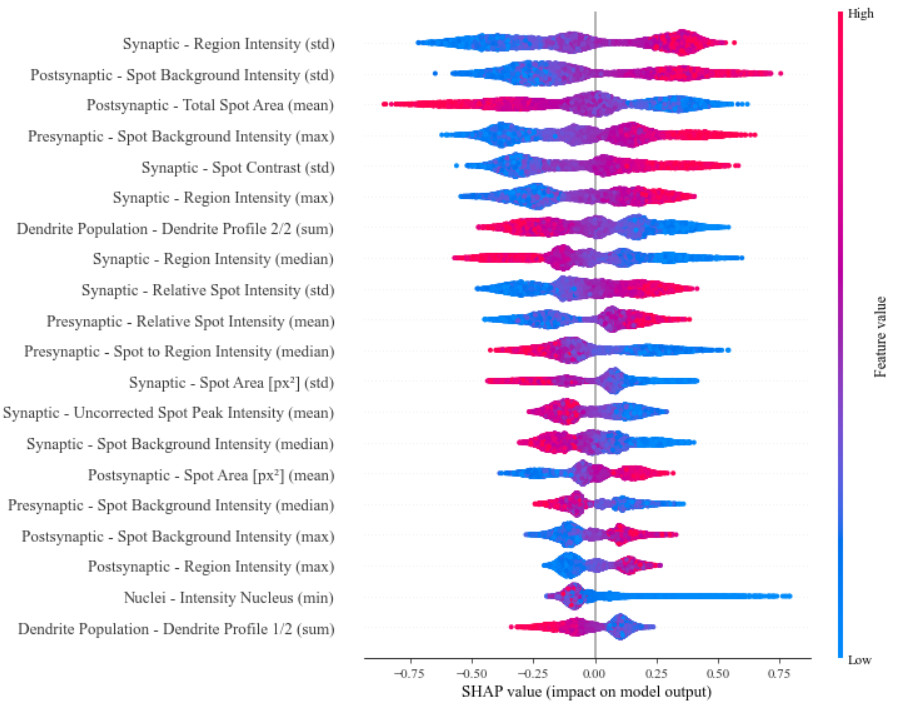

Figure S11. 20 most important features for the XGBoost model measured by SHAP values, dose 30. The y-axis indicates the cell feature names sorted by the importance. The x-axis is the SHAP values of each feature. The color represents the feature value (red high, blue low).
